# Characterization of a high-intensity band that cross-reacts with FLAG-M2 antibodies in immunoblots in a subset of laboratory strains of Saccharomyces cerevisiae

**DOI:** 10.1101/2025.04.15.648968

**Authors:** Nicolle K. Reid, Amy C. Liang, Mara C. Duncan

## Abstract

Epitope tags are commonly used for various purposes in research labs. The DYKDDDDK-peptide epitope, trademarked as the FLAG epitope, is a commonly used epitope tag. It is often used for monitoring protein levels and for affinity chromatography. Multiple DYKDDDDK-binding antibodies are available; however, the mouse monoclonal anti-FLAG M2 is widely used due to its commercial availability in several formats. Many laboratory *Saccharomyces cerevisiae* strains, including the BY4741 strain that was used in multiple systematic deletion and tagging libraries, have a high-intensity band that cross-reacts with the FLAG-M2 antibody. The presence of this high-intensity cross-reactive band can be problematic in some applications. Here, we show that despite high-intensity in immunoblots, the cross-reacting band is not enriched by FLAG-M2 affinity beads under native conditions. We also report the fortuitous identification of a strain closely related to BY4741 that lacks the high-intensity cross-reactive band. Finally, contrary to anecdotal reports, we determined that the high-intensity cross-reacting band is not Rtf1. These findings and resources should assist other researchers using the DYKDDDDK-epitope for immunoblots and affinity chromatography.

## Introduction

Epitope tags are small protein modules that can be added to proteins using molecular genetics approaches that can be recognized with high affinity antibodies (1, 2). Epitope tagging has become ubiquitous in cell biological studies because it removes the need to generate protein-specific antibodies, which is expensive, time-consuming, and does not always yield antibodies suitable for desired applications. Moreover, using any antibody carries the risk of cross-reactivity with endogenous proteins, which must be carefully assessed for each new antibody, organism or cell type, and application.

The FLAG tag is a commonly used epitope tag. It is a short artificial peptide with the sequence DYKDDDDK(3, 4). It is often used in 1-5 copies appended to the N or C-terminus of a protein. Its small size makes it suitable for tandem tagging approaches (5-7). Multiple commercial antibodies recognize the FLAG tag. One commonly used antibody is the FLAG-M2 mouse monoclonal antibody, which is suitable for immunoblot and affinity approaches (8). Numerous commercial products are available using FLAG-M2 for both applications.

Although FLAG-M2 has low cross-reactivity in many samples, cross-reactivity has been previously described in some systems (9-11). Cross-reactive bands can be helpful by acting as a loading control band; however, if the cross-reactive band is the same or near the molecular weight of a protein of interest, this can be problematic. Moreover, if the cross-reactive band binds to affinity reagents, it can reduce the overall yield of affinity protocols or complicate the interpretations of experimental outcomes with affinity reagents.

Over the course of our studies, we noticed a band with strong cross-reactivity for FLAG-M2 at approximately 95kDa in cell lysates from cells derived from the BY4741 *Saccharomyces cerevisiae* strain. This cross-reactive band is intense and can easily obscure the signal of proteins that migrate at or near 95 kDa. However, some of our strains lacked this band, revealing unexpected strain-to-strain variability in the presence of this epitope. Here, we report that the high-intensity cross-reactive band is found in the S288C background strains, BY4741 and BY4742, which were used to make several systematic libraries, and in additional S288C strains (12-14). However, the band is much less intense in the S288C background strains FY4, FY5, and DYB1034. In addition, we show that the high-intensity cross-reactive band is not enriched by two affinity chromatography resins, suggesting that even strains containing the high-intensity cross-reactive band are suitable for use with FLAG-binding affinity approaches. Finally, we explore anecdotal reports that the high-intensity cross-reacting band is Rtf1 and demonstrate that Rtf1 is not the high-intensity cross-reacting band. These data provide valuable insight into the potentially confounding FLAG-M2 cross-reactive band observed in some yeast strains.

## Results

### The FLAG-M2 antibody detects a high-intensity band in lysates of BY4741 but not FY4 or FY5

We observe a high-intensity ∼95 kDa band in lysates of BY4741 and BY4742 and their direct progeny when probed with the FLAG-M2 antibody (Fig 1a). In fluorescent western blots run in the linear range, the cross-reacting band is stronger than some endogenously tagged proteins carrying 3 or 5 copies of the FLAG epitope.

**Figure 1.**
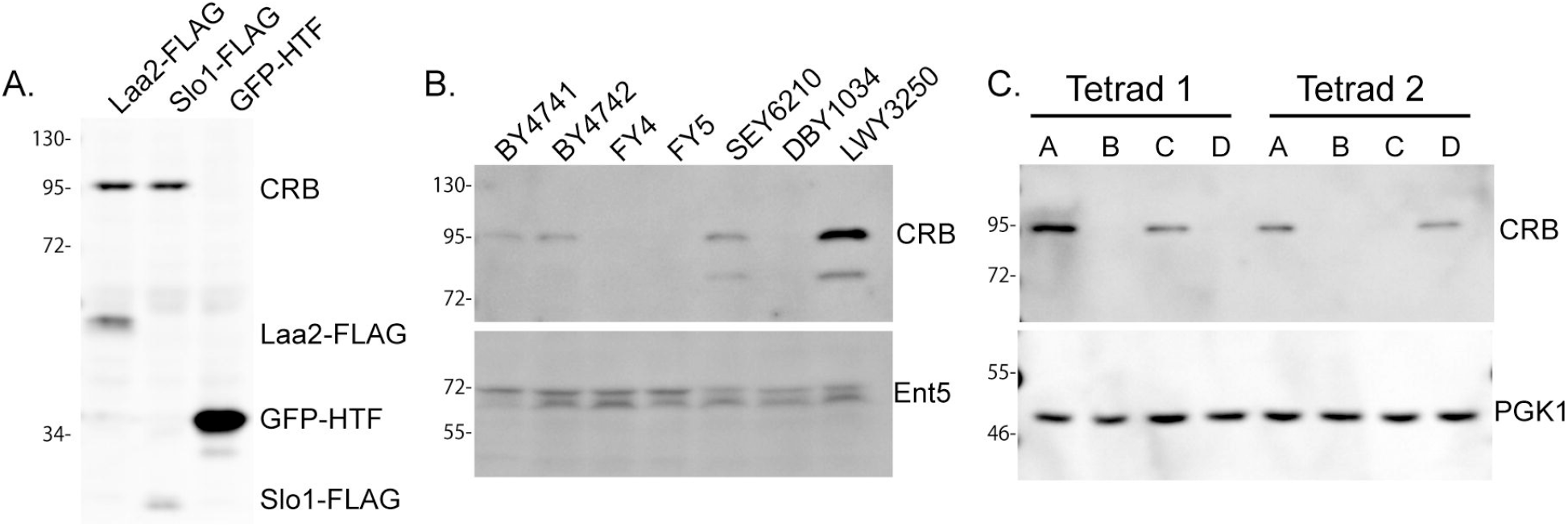
Lysates from progeny ofBY4741 contain a 95kDa band that cross-reacts with the Flag-M2 antibody in immunoblots. A. Lysates from strains derived from BY4741 (Laa2-FLAG, Slol -FLAG) and or crosses between progeny ofBY4741 and FY4 (GFP-HIS-TEV-FLAG(HTF)) expressing the indicated FLAG tagged proteins probed with the FLAG-M2 antibody. Cross-reacting band is labeled CRB. B. Lysates from indicated wild-type yeast strains probed with FLAG-M2 or anti-sera that recognizes Ent5. C. Lysates from two tetrads ofa cross between BY4741 and FY5 probed with FLAG-M2 antibody or a PGKl antibody.

Curiously, in some of the strains generated in our lab, the 95 kDa band is notably less intense (Fig 1A). We determined that the strains with the less intense band derived from crosses of descendants of BY4741 or BY472 and FY4 or FY5. FY4 and FY5 are prototrophic strains considered closely related to BY4741. Indeed, FY4 was used in the final cross that produced BY4741(15).

To confirm that the less intense band in our strains originates from FY4 and/or FY5, we performed immunoblots on multiple wild-type laboratory yeast strains (Fig 1b). The high-intensity band was observed in the S288C background strains BY4741, BY4742, SEY6210, and LWY3250, however, it was far less intense in lysates from FY4 and FY5, as well as in DBY1034, another S288C wild-type strain (16-19). We note the band was slightly more intense in LWY3250 than in BY4741, BY4742, or SEY6210. These results show that the intensity of this band is highly variable in laboratory yeast strains.

To determine if the different intensity of the 95 kDa band in different S288C strains is due to a single genetic locus, we crossed FY5 versus BY4741 and performed western blots on the progeny from multiple tetrads. We observed that the high-intensity band segregated 2:2, meaning that two spores from each tetrad carried the high-intensity band, and two carried the low-intensity band (Fig 1c). This indicates that a single genetic locus likely causes the high-intensity 95 kDa band.

### The 95 kDa cross-reacting band does not bind FLAG-M2 or L5-rat monoclonal affinity beads

One notable concern about any cross-reacting band is that it may compete with FLAG-tagged proteins for binding to FLAG-M2 affinity reagents. This competition could reduce the binding capacity of the affinity reagent for the desired tagged protein and raise the possibility that the cross-reacting band or its interacting partners would complicate downstream applications or interpretations. To determine if the 95 kDa band binds to FLAG-M2 affinity reagents, we performed immunoprecipitations from cells containing the high-intensity band and other FLAG-tagged proteins. We found that the 95 kDa band was not enriched in immunoprecipitations using FLAG-M2 resin (Fig 2a). We found similar results using a second resin that uses the L5 rat monoclonal antibody, which recognizes the DYKDDDDK-peptide (Fig 2b)(20). This suggests that although the 95 kDa band is recognized by FLAG-M2 under the denaturing conditions used for immunoblots, it is not recognized by either FLAG-M2 or L5 antibodies under the non-denaturing conditions we use for affinity purification.

**Figure 2.**
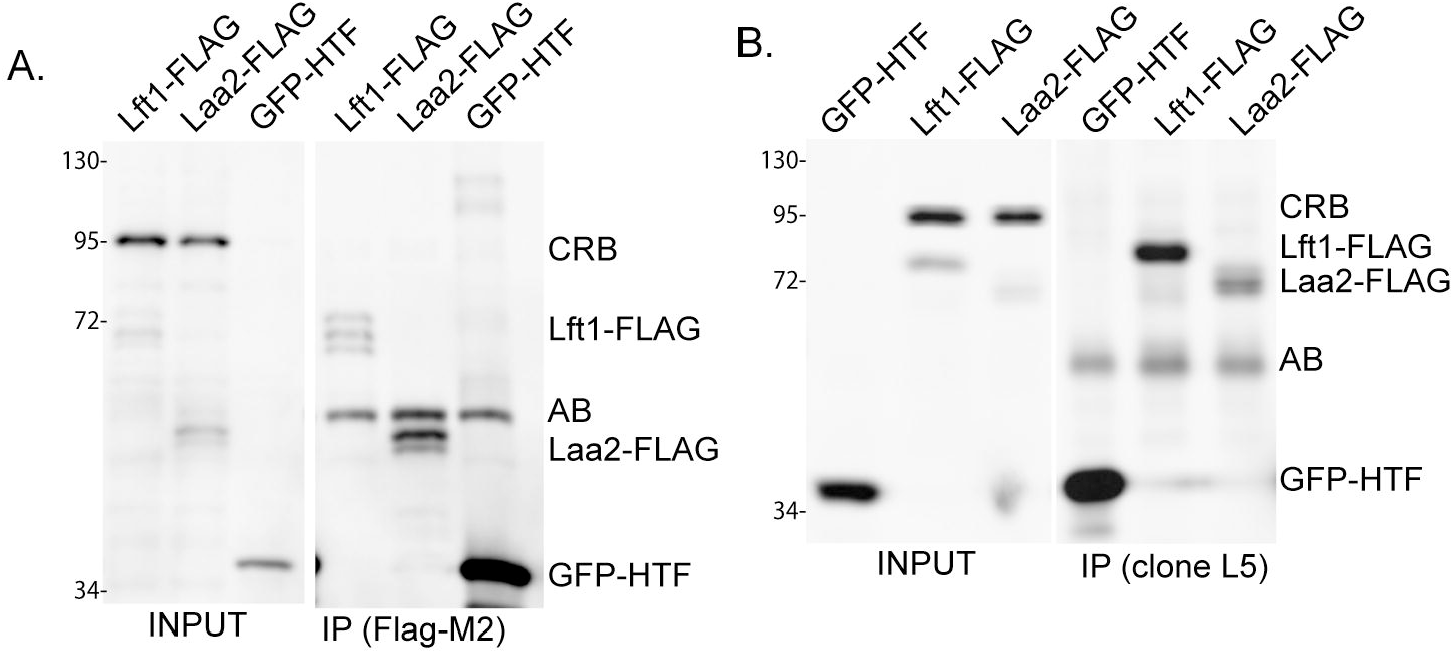
The 95 kDa cross-reacting band is not enriched with DYKDDDDK affinity reagents under conditions used for immuno-precipitation. A. Lysates from cells expressing indicated FLAG-tagged proteins were subjected to immuno-precipitation (IP) using FLAG-M2 conjugated agarose beads, eluted and probed with FLAG-M2 antibody. AB indicates the FLAG-M2 antibody band recognized by the secondary antibody. B Lysates expressing indicated FLAG-tagged proteins were subjected to immuno precipitation using anti-DYKDDDDK-L5 antibody conjugated magnetic beads, eluted, and probed with FLAG-M2 antibody. AB indicates the L5 antibody band recognized by the secondary antibody.

### The 95kDa band is not Rtf1

The 95kDa band was proposed to be Rtf1 based on reports of the enrichment of the Rtf1-containing PAF complex on Flag-M2 beads, the presence of a region in Rtf1(^196^DYKDDEGS) with notable similarity to the DYKDDDDK peptide, and the similarity in size between the 95kDa band and Rtf1 (Karen Arndt, personal communication and (21)). To examine this possibility, we probed lysates from three different rtf1Δ strains with FLAG-M2 antibody and an antibody that recognizes endogenous Rtf1 (Fig 3). Although the high-intensity FLAG-M2 cross-reacting band was absent from two *RTF1* delete strains, it was present in a third *RTF1* delete strain tested. Moreover, the Rtf1 band migrates faster than the 95kDa band (Fig 3). Together, these results indicate that the 95kDa band is not Rtf1.

**Figure 3.**
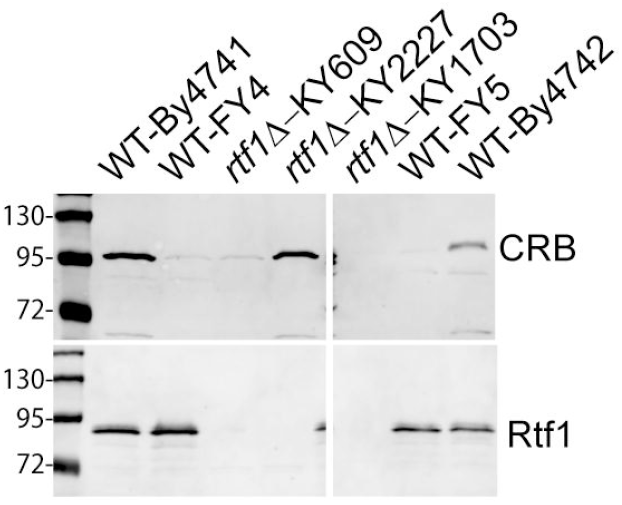
The high-intensity 95 kDa cross-reacting band is not Rtfl. Lysates from indicated strains were subjected to immunoblot with FLAG-M2 or antisera that recognizes Rtfl. Molecular weight markers are shown as a reference for the faster migration of Rtfl compared to CRB.

## Discussion

Here, we report that the intensity of a FLAG-M2 cross-reacting band is variable even in highly related yeast strains. Because strains derived from BY4741, FY4, SEY6210, and DBY1034 are commonly used in research labs, other labs may encounter unexpected differences in the FLAG-M2 cross-reacting band in strains considered isogeneic. Our results should provide peace of mind to researchers concerned that their strains have acquired an unknown FLAG-tagged protein.

We determined that this intensity difference maps to a single genetic locus and that it is not Rtf1. It remains to be determined what the locus is and whether it alters the expression level of a protein containing the cross-reacting peptide or directly impacts the sequence of the cross-reacting peptide. Future work using pooled segregant sequencing from the backcrossed strains used in Fig1C should be able to identify the relevant genetic difference (22).

We also report that the FLAG-M2 cross-reacting band is not enriched by FLAG-M2 or anti-DYKDDDDK L5 rat monoclonal antibodies under our tested conditions. Notably, we have only used a small number of buffer conditions. Therefore, if competition for the affinity reagent or contamination from the cross-reacting band is a concern for other researchers, a pilot study should be performed to determine the extent to which the 95 kDa band binds under the affinity conditions used.

We showed that a single backcross to FY4 or FY5 is sufficient to generate a replacement for BY4741. This can provide an option for researchers needing to visualize proteins whose signal would be obscured by the high-intensity band. We note, however, that the high-intensity cross-reacting band can be useful as an internal loading control. Therefore, unless the band causes a problem for immunoblotting, it may be preferable to retain the cross-reacting band.

In summary, this study reveals insight into a potentially vexing 95 kDa band that cross-reacts strongly with the FLAG-M2 antibody in immunoblots of lysates from some Saccharomyces cerevisiae strains. Our characterization of its absence in some strains closely related to BY4741 and its absence in affinity purification reactions under native conditions should provide valuable insight to other researchers who are confused or concerned by the presence of the band.

## Materials and Methods

The strains and oligonucleotides used for this study are listed in Tables 1 and 2 (15-19, 23). Tags were integrated into the genome using a one-step PCR-based method (24, 25). Yeast cells were grown in yeast/peptone medium supplemented with 2% glucose and a mixture of adenine, uracil, and tryptophan (YPD+AUT).

FlagM2 antibody (RRID:AB_439685) was from Sigma. PGK1 antibody (RRID:AB_2532235) was from Thermo Fisher Scientific. Ent5 antibodies were described previously(26). Rtf1 antisera was a gift from Karen Arndt. Alexa Fluor 647-conjugated goat anti-mouse (RRID:AB_141698) and Alexa Fluor 647 goat anti-rabbit (RRID:AB_141663) antibody were from Life Technologies.

For immunoblotting, after SDS-PAGE, samples were transferred to nitrocellulose, blocked with 5% nonfat dry milk in TBS-T (137 mm NaCl, 15.2 mm Tris-HCl, 4.54 mm Tris, 0.896 mm Tween 20), and then probed with primary and fluorescent secondary antibodies. Fluorescence signals were detected on an Azure 600 imaging system (Azure Biosciences).

For immunoblotting without immunoprecipitation, lysates were generated by glass-bead lysis in hot laemmli sample buffer. For immunoprecipitations, lysates were generated by glass-bead lysis in HEKG5 with 0.3% CHAPS (20 mm HEPES, pH 7.5, 1 mm EDTA, 150 mm KCl, 5% glycerol) with protease inhibitor mixture without EDTA (Promega). The lysates were clarified by centrifugation at 13,000 rpm for 10 min at 4 °C. Protein concentrations were determined using Biorad Protein Assay (Biorad), and protein concentrations were adjusted in HEKG5 with 0.3% CHAPS. For immunoprecipitation, 6 μls of EZview™ Red ANTI-FLAG® M2 Affinity Gel (Sigma) were incubated with 2mg of lysate in 500 μl reactions or 50 μl of Anti-DYKDDDDK Magnetic Agarose (Pierce) were incubated with 10mg of lysate in 10 ml reactions. The beads were washed three times with ice-cold HEKG5 with 0.3% CHAPS, and the bound proteins were eluted with laemmli sample buffer.

## Acknowledgements

We thank Karen Arndt for useful discussions and generous sharing of reagents. This work was supported by the National Institutes of Health (R01-GM129255, UL1-TR002240) and used information available through the yeast genome database, which is supported by U24-HG001315, U41-HG002273, and U24-HG010859. AL and NR were supported by the federal work-study program through the Department of Education.

